# Adaptation mechanisms of *Listeria monocytogenes* to quaternary ammonium compounds

**DOI:** 10.1101/2023.03.30.534860

**Authors:** Lisa Maria Schulz, Fabienne Dreier, Lisa Marie de Sousa Miranda, Jeanine Rismondo

## Abstract

*Listeria monocytogenes* is ubiquitously found in nature and can easily enter food-processing facilities due to contaminations of raw materials. Several countermeasures are used to combat contamination of food products, for instance the use of disinfectants that contain quaternary ammonium compounds, such as benzalkonium chloride (BAC) and cetyltrimethylammonium bromide (CTAB). In this study, we assessed the potential of the commonly used wildtype strain EGD-e to adapt to BAC and CTAB under laboratory growth conditions. All BAC-tolerant suppressors exclusively carried mutations in *fepR* or its promoter region likely resulting in the overproduction of the efflux pump FepA. In contrast, CTAB- tolerance was associated with mutations in *sugR*, which regulates the expression of the efflux pumps SugE1 and SugE2. *L. monocytogenes* strains lacking either FepA or SugE1/2 could still acquire tolerance towards BAC and CTAB. Genomic analysis revealed that the overproduction of the remaining efflux system could compensate for the deleted one. Even in the absence of both efflux systems, tolerant strains could be isolated, which all carried mutations in the diacylglycerol kinase encoding gene *lmo1753* (*dgkB*). DgkB converts diacylglycerol to phosphatidic acid, which is subsequently re-used for the synthesis of phospholipids suggesting that alterations in membrane composition could be the third adaptation mechanism.

**Originality-Significance Statement:** Survival and proliferation of *Listeria monocytogenes* in the food industry is an ongoing concern, and while there are various countermeasures to combat contamination of food products, the pathogen still successfully manages to withstand the harsh conditions present in food-processing facilities, resulting in reoccurring outbreaks, subsequent infection, and disease. To counteract the spread of *L. monocytogenes* it is crucial to understand and elucidate the underlying mechanism that permit their successful evasion. We here present various adaptation mechanisms of *L. monocytogenes* to withstand two important quaternary ammonium compounds.

## Introduction

*Listeria monocytogenes* is one of the most successful foodborne pathogens worldwide. In high-risk groups such as immunocompromised individuals, the elderly, or pregnant women, an infection can cause invasive listeriosis, resulting in a hospitalization rate of ∼95% and a fatality rate of ∼13% (Allerberger and Wagner 2010; EFSA and ECDC 13/12/2022). Due to its ability to persist in a wide range of environmental stresses found in the food-processing industry, infection of *L. monocytogenes* is often associated with ingestion of contaminated ready-to-eat foods, such as ice cream (Rietberg *et al*. 2016), cheese (Carlin *et al*. 2022), or processed meat (Thomas *et al*. 2020). The pathogen does not only pose a threat due to its ability to withstand common stresses such as extreme temperatures, pH, or high salt concentrations (up to 20%), but also due to its potential to adapt to biocides that are commonly found in disinfectants or sanitizers used in food-processing plants (Osek *et al*. 2022). The most frequently used antimicrobial components in disinfectants are a mixture of quaternary ammonium compounds (QACs) that are characterized by its ammonium ion linked to either an alkyl or aryl group. The chain length of QACs determines the antimicrobial potency, hence, a length of C14 or C16 are ideally used against Gram-positive and Gram-negative bacteria, respectively (Zinchenko *et al*. 2004). *L. monocytogenes* strains with decreased sensitivity to QACs have been isolated from different locations around the world (Aase *et al*. 2000; Romanova *et al*. 2006; Romanova *et al*. 2002; Soumet *et al*. 2005; Meier *et al*. 2017; Lundén *et al*. 2003; To *et al*. 2002). This adaptation frequently resulted in cross-adaptation to other disinfectants and antimicrobial agents such as gentamycin, kanamycin or ciprofloxacin (Aase *et al*. 2000; Romanova *et al*. 2006; Lundén *et al*. 2003; Guérin *et al*. 2021; Guérin *et al*. 2014; Rakic-Martinez *et al*. 2011; Bland *et al*. 2022), emphasising the importance of elucidating the underlying genetic basis. Two of the most extensively used QACs are benzalkonium chloride (BAC) and cetyltrimethylammonium bromide (CTAB) (Jiang *et al*. 2016; Martínez-Suárez *et al*. 2016). While a variety of different mechanisms have been linked to increased tolerance towards BAC in isolated *L. monocytogenes* strains, as far as we know, no previous research has investigated the underlying genetics of CTAB tolerance, which is often merely mentioned in association with cross-adaptation towards BAC (Jiang *et al*. 2020; Müller *et al*. 2013; Müller *et al*. 2014).

Several factors have been identified in *L. monocytogenes* strains isolated from food-processing facilities that are associated with the presence or overexpression of efflux systems that aid in extruding the toxic compounds from the intracellular space. Determinants for BAC tolerance are often found on genetic elements, and include genes encoding efflux pumps such as *qacH*, located on the transposon Tn*6188* (Müller *et al*. 2013; Müller *et al*. 2014), *emrE*, located on the genomic island LGI1 (Kovacevic *et al*. 2015), or *bcrABC*, which encodes the TetR-family transcriptional regulator BcrA and two small multidrug resistance (SMR) efflux pumps, BcrB and BcrC (Elhanafi *et al*. 2010; Dutta *et al*. 2013; Minarovičová *et al*. 2018). The *sugRE1E2* operon (short: *sug* operon) was analysed as a chromosomal counterpart of *bcrABC* in the laboratory wildtype strain EGD-e and was correspondingly found to be important for tolerance towards QACs such as BAC and CTAB. The genes of the *sug* operon code for the TetR-family regulator SugR and the two SMR efflux pumps SugE1 and SugE2. The self-repressor SugR negatively regulates the operon in the absence of BAC and accordingly, both SugE1 and SugE2 showed increased expression in the presence of the QAC (Jiang *et al*. 2020). In addition, several studies showed increased expression of the major facilitator superfamily transporter MdrL in isolated, as well as, BAC-adapted *L. monocytogenes* strains, suggesting a direct contribution of this transporter to BAC tolerance (Romanova *et al*. 2006; Yu *et al*. 2018). However, a clean deletion of the transporter in EGD-e only resulted in a growth defect in the presence of BAC, but no change in the minimal inhibitory concentration (MIC). Additionally, its role in export of cefotaxime and EtBr, that was previously described, could not be confirmed for the laboratory model strain (Mata *et al*. 2000; Jiang *et al*. 2019). It has to be mentioned that the tolerance in isolated or adapted *L. monocytogenes* strains was often transient and lost after passaging the strains in the absence of QAC stress. In contrast, the chromosomally encoded multidrug and toxic compound (MATE) efflux pump FepA was recently identified as the dominant, stable mode of tolerance for BAC in a screen of over 60 serial adapted produce-associated *L. monocytogenes* and other *Listeria* spp. strains (Bolten *et al*. 2022). An independent study of biocide-adapted strains further supported these findings by showing that 94% of the adapted strains possessed mutations in the gene coding for the TetR-like transcriptional regulator FepR that was previously shown to repress its own expression and the expression of the efflux pump encoding gene *fepA* (Douarre *et al*. 2022). The identified mutations in *fepR* were thus proposed to increase expression of FepA, resulting in enhanced tolerance towards BAC, as well as norfloxacin and ciprofloxacin (Guérin *et al*. 2014).

Here we show that the laboratory wildtype strain EGD-e readily acquires stable mutations in the transcriptional regulator *fepR* and that the successional overexpression of the efflux pump FepA is responsible for the increased tolerance towards BAC, CTAB, ciprofloxacin and gentamycin. We further successfully evolved suppressors in presence of CTAB which exclusively carried mutations in the TetR-like transcriptional regulator *sugR* resulting in the overexpression of the SMR efflux pumps SugE1 and SugE2. *L. monocytogenes* strains lacking either *fepA* or *sugE1*/*2* could still acquire tolerance towards CTAB and BAC by overexpressing the remaining efflux system. In addition, we could further evolve BAC- and CTAB-tolerant strains in the absence of the two major QAC efflux systems, which acquired mutations in a putative diacylglycerol kinase.

## Materials and Methods

### Bacterial strains and growth conditions

All strains and plasmids used in this study are listed in Table S1. *Escherichia coli* strains were grown in lysogeny broth (LB) medium and *L. monocytogenes* strains in brain heart infusion (BHI) medium at 37°C unless otherwise stated. If appropriate, antibiotics and supplements were added to the medium at the following concentrations: for *E. coli* cultures ampicillin (Amp) at 100 µg ml^-1^, kanamycin (Kan) at 50 µg ml^-1^, and for *L. monocytogenes* strains, chloramphenicol (Cam) at 7.5 µg ml^-1^, Kan at 50 µg ml^-1^, erythromycin (Erm) at 5 µg ml^-1^ and IPTG at 1 mM.

### Strain and plasmid construction

All primers used in this study are listed in Table S2. For the markerless in-frame deletion of *fepA* (*lmo2087*) and *sugE1/2* (*lmo0853-lmo0854*), approximately 1-kb DNA fragments up- and downstream of the *fepA* gene were amplified by PCR using the primer pairs LMS484/LMS485 and LMS486/LMS487 (*fepA*) and JR247/JR248 and JR249/JR250 (*sugE1/2*). The resulting PCR products were fused in a second PCR using primers LMS485/LMS487 (*fepA*) and JR247/JR250 (*sugE1/2*). The products were cut with *Kpn*I and *Sal*I and ligated into pKSV7 that had been cut with the same enzymes. The resulting plasmids pKSV7-Δ*fepA* and pKSV7-Δ*sugE1/2* were recovered in *E. coli* XL1-Blue, yielding strains EJR230 and EJR229, respectively. Plasmids pKSV7-Δ*fepA* and pKSV7- Δ*sugE1/2* were transformed into *L. monocytogenes* EGD-e and the genes deleted by allelic exchange according to a previously published method (Camilli *et al*. 1993) yielding strains EGD-e Δ*fepA* (LJR261) and EGD-e Δ*sugE1/2* (LJR262), respectively. For the construction of the Δ*fepA*Δ*sugE1/2* double deletion strain, plasmid pKSV7-Δ*sugE1/2* was transformed into EGD-e Δ*fepA* and *sugE1/2* was deleted by allelic exchange, resulting in strain LJR329. For the construction of pIMK3-*fepA* and pIMK3-*sugE1/2*, the *fepA* and *sugE1/2* genes were amplified using the primer pairs LMS478/LMS479 and JR262/JR263, respectively. Fragments were cut with enzymes *Nco*I and *Sal*I and ligated into plasmid pIMK3 that had been cut with the same enzymes. The resulting plasmids pIMK3-*fepA* and pIMK3-*sugE1/2* were recovered in *E. coli* XL10-Gold yielding strains EJR227 and EJR259, respectively. Both plasmids were transformed into *L. monocytogenes* strain EGD-e, resulting in the construction of strains LJR231 and LJR301, respectively, in which the expression of *fepA* and *sugE1/2* is under the control of an IPTG- inducible promoter. For the construction of a *fepA* complementation strain, plasmid pIMK3-*fepA* was transformed into EGD-e Δ*fepA* yielding strain LJR265.

For the construction of pWH844-*fepA*, the *fepA* gene was amplified with primer pairs FD1/FD2 using EGD-e wild type DNA or the DNA of suppressor strain LJR218 as template. The PCR fragments were digested with *Bam*HI and *Sal*I and ligated into pWH844 that had been cut with the same enzymes. The resulting plasmids pWH844-*fepR^WT^* and pWH844-*fepR^L24F^* were recovered in XL10-Gold, yielding strains EJR242 and EJR248, respectively.

For the construction of promoter *lacZ* fusions, the promoter region of *fepR* was amplified with primers FD5 and FD6 using genomic DNA of the *L. monocytogenes* wildtype strain EGD-e or the BAC-tolerant strains EGD-e *P_fepR_^G-27T^* (LJR188) or EGD-e *P_*fepR*_^A-33G^* (LJR215) as template DNA. The PCR fragments were digested with *Bam*HI and *Sal*I and ligated into plasmid pPL3e-*lacZ*, which contains the promoter-less *lacZ* gene. The resulting plasmids pPL3e-*P_fepR_-lacZ*, pPL3e-*P_fepR_^G-27T^-lacZ* and pPL3e-*P_fepR_^A-33G^-lacZ* were recovered in *E. coli* DH5α yielding strains EJR257, EJR258 and EJR260, respectively. Plasmids pPL3e- *P_fepR_-lacZ*, pPL3e-*P_fepR_^G-27T^-lacZ* and pPL3e-*P_fepR_^A-33G^-lacZ* were subsequently transformed into EGD-eyielding strains EGD-e pPL3e-*P*_*fepR*_-*lacZ* (LJR336), EGD-e pPL3e-*P*_*fepR*_^*G-27T*^-*lacZ* (LJR302) and EGD-e pPL3e-*P*_*fepR*_^*A-33G*^-*lacZ* (LJR303).

### Generation of suppressors and whole genome sequencing

For the generation of BAC-adapted suppressors, stationary or exponentially grown EGD-e cultures were selected on BHI plates containing 4 µg ml^-1^ or 6 µg ml^-1^ BAC. For the stationary grown EGD-e cultures, overnight cultures were adjusted to an OD_600_ of 0.1 and 100 µl were plated on BHI plates containing 4 µg ml^-1^ BAC. For exponentially grown cultures, overnight cultures of EGD-e were adjusted to an OD_600_ of 0.1 and grown until they reached an OD_600_ of 0.3-0.5. Cultures were then adjusted to an OD_600_ of 0.1 and 100 µl were plated on BHI plates containing 4 µg ml^-1^ and 6 µg ml^-1^ BAC. The plates were incubated at 37°C overnight and single colonies were re-streaked twice on 4 µg ml^-1^ and 6 µg ml^-1^ BAC, respectively. For adaptation of the wildtype strain to CTAB, as well as EGD-e Δ*fepA*, EGD-e Δ*sugE1/2* and EGD-e Δ*fepA* Δ*sugE1/2* to BAC and CTAB, overnight cultures of the different strains were adjusted to an OD_600_ of 0.1 and grown to an OD_600_ of 0.3-0.5. Cultures were adjusted again to an OD_600_ of 0.1 and 100 µl were plated on BHI plates supplemented with 4 µg ml^-1^, 5 µg ml^-1^ or 6 µg ml^-1^ BAC and 2 µg ml^-1^ or 4 µg ml^-1^ CTAB. Plates were incubated at 37°C overnight or in the case of EGD-e Δ*fepA* and EGD-e Δ*fepA* Δ*sugE1/2* in presence of BAC for 2 days. Again, single colonies were re-streaked twice on BHI plates supplemented with the selective pressure they were originally isolated from. Genomic DNA of a selection of BAC- and CTAB- adapted strains was isolated and either pre-screened for mutations in *fepR* or *sugR*, or send to SeqCoast Genomics (Portsmouth, NH, United States) for whole genome sequencing (WGS). The genome sequences were determined by short read sequencing (150-bp paired end) using an Illumina MiSeq system (San Diego, CA, United States). The reads were trimmed and mapped to the *L. monocytogenes* EGD-e reference genome (NC_003210) using the Geneious prime^®^ v.2021.0.1 (Biomatters Ltd., New Zealand). Single nucleotide polymorphisms (SNPs) with a variant frequency of at least 90% and a coverage of more than 25 reads were considered as mutations. All identified mutations were verified by PCR amplification and Sanger sequencing.

### Drop dilution assay

Overnight cultures of the indicated *L. monocytogenes* strains were adjusted to an OD_600_ of 1. IPTG and Kan were supplemented to the overnight cultures of strains carrying pIMK3- plasmids. 5 µl of serial dilutions of each culture were spotted on BHI agar plates, BHI agar plates containing 4 µg ml^-1^ BAC, 6 µg ml^-1^ BAC, 2 µg ml^-1^ CTAB, 4 µg ml^-1^ CTAB, 1 µg ml^-1^ ciprofloxacin, 0.5 µg ml^-1^ gentamycin, or 1 µg ml^-1^ cefuroxime. Where indicated, plates were supplemented with 1 mM IPTG. Images of plates were taken after 20-24 h of incubation at 37°C. Drop dilution assays were repeated at least three times.

### Ethidium bromide assay

The ethidium bromide assay was performed as previously described with minor modifications (Kaval *et al*. 2015). Briefly, overnight cultures of the indicated *L. monocytogenes* strains were diluted to an OD_600_ of 0.05 in fresh BHI medium and grown until an OD_600_ of 0.4-0.6. Cells of 2 ml were harvested by centrifugation at 1,200 x g for 5 min, washed once in 1 ml PBS buffer (pH 7.4) and finally re-suspended in 1 ml PBS buffer (pH 7.4). Next, the OD_600_ of each sample was adjusted to 0.3 in PBS (pH 7.4) and 180 µl transferred into the wells of a black 96-well plate. 20 µl of 50 µg ml^-1^ ethidium bromide was added to each well and the absorbance was measured using the Synergy^TM^ Mx microplate reader (BioTek) at 500 nm excitation and 580 nm emission wavelengths for 50 min.

### Expression and purification of His-FepR

For the overexpression of His-FepR and His-FepR^L24F^, plasmids pWH844-*fepR* and pWH844-*fepR^L24F^* were transformed into *E. coli* strain BL21 and the resulting strains grown in LB broth supplemented with Amp at 37°C. At an OD_600_ of 0.6-0.8, the expression of *his*-*fepR* and *his*-*fepR^L24F^* was induced by the addition of 1 mM IPTG and the strains were grown for another 2 h at 37°C. Cells were collected by centrifugation at 11,325 x g for 10 min, washed once with 1x ZAP buffer (50 mM Tris-HCl, pH 7.5, 200 mM NaCl) and the cell pellet stored at −20°C until further use. The cell pellets were re-suspended in 1x ZAP buffer and cells passaged three times (18,000 lb/in2) through an HTU DIGI-F press (G. Heinemann, Germany). The cell debris was subsequently collected by centrifugation at 46,400 x g for 30 min. The supernatant was subjected to a Ni^2+^ nitrilotriacetic acid column (IBA, Göttingen, Germany) and His-FepA and His-FepA^L24F^ were eluted using an imidazole gradient. Elution fractions were analysed by SDS-PAGE and selected fractions subsequently dialysed against 1x ZAP buffer with a spatula pinch of EDTA at 4°C overnight. Protein concentrations were determined by a Bradford protein assay (Bradford 1976) using the Bio-Rad protein assay dye reagent concentrate. Bovine serum albumin was used for a standard curve. The protein samples were stored at 4°C until further use. Two independent purifications were performed for each protein.

### Electrophoretic mobility shift assay (EMSA)

EMSAs were performed as described elsewhere with minor modifications (Dhiman *et al*. 2014). Briefly, a 150 bp-DNA fragment containing the *fepR* promoter was amplified using primers FD3 and FD4 from genomic DNA isolated from the wildtype strain EGD-e or EGD-e *P_fepR_^G-27T^* (LJR188) and EGD-e *P_*fepR*_^A-33G^* (LJR215). For the comparison of the binding abilities of His-FepR and His-FepR^L24F^ to the *fepR* promoter, 25, 50 and 100 pmol of each protein were mixed with 250 pmol *fepR* promotor DNA. To compare the binding ability of His-FepR to the wildtype and the mutated *fepR* promoters, 25, 50 and 100 pmol of His-FepR were mixed with 250 pmol DNA of either the wildtype or the mutated *fepR* promoters. Apart from DNA and protein, 20 µl binding reactions contained 1 µl of DNA loading dye (50% glycerol, 0.1% bromophenol blue, 1x TAE, H_2_O), 50 mM NaCl, 2 µl 10x Tris-acetate buffer (250 mM Tris-base in H_2_O, set to pH 5.5 with acetic acid), 0.15 mM bovine serum albumin, 2.5 mM EDTA, 10% glycerol and 20 mM DTT. The samples were incubated for 5 min at 25°C and subsequently separated on 8% native Tris-acetate gels (6% polyacrylic acid, 1x Tris-acetate buffer, 0.15% ammonium persulfate, 0.83% tetramethylethylenediamine) in 0.5x TBE-buffer (0.5 M Tris base, 0.5 M boric acid, 1 mM Na_2_EDTA, pH 10). A pre-run of the gels was performed for 90 min at 50 V before the samples were loaded. The run was performed at 50 V for 2.5 h. The gels were stained in 50 ml 0.5x TBE-buffer containing 5 µl HDGreen^TM^ Plus DNA dye (INTAS, Göttingen, Germany) for 5 min and washed for 5 min with 0.5x TBE-buffer, rinsed three times with water and then washed with water for 30 min. The DNA bands were visualized using a Gel Doc^TM^ XR+ (Bio Rad, Munich, Germany).

### β-galactosidase assays

For the comparison of the activity of the wildtype and mutated *fepR* promoters, overnight cultures (supplemented with Erm) of the indicated *L. monocytogenes* strains were diluted to an OD_600_ of 0.05 in BHI medium and grown for 5 h at 37°C. To determine the response of the *fepR* promoter to BAC, overnight cultures of EGD-e *P_fepR_-lacZ* were diluted to an OD_600_ of 0.05 in BHI medium and grown at 37°C to an OD_600_ of 0.5 (± 0.05). The culture was divided into two flasks and incubated for an additional 2 h at 37°C in the presence of 2.5 µg ml^-1^ BAC or in the absence of BAC. The final OD_600_ was measured for all cultures prior sample collection. For both assays, 1 ml of the corresponding cultures were collected, re-suspended in 100 µl ABT buffer (60 mM K_2_HPO_4_, 40 mM KH_2_PO_4_, 100 mM NaCl, 0.1% Triton X-100; pH 7), snap-frozen in liquid nitrogen and stored at −80°C until further use. The sample preparation was performed as described previously (Gründling *et al*. 2004; Rismondo *et al*. 2019). Briefly, samples were thawed, and 10-fold dilutions were prepared in ABT buffer. 50 µl of each dilution were mixed with 10 µl of 0.4 mg ml^-1^ 4-methyl-umbelliferyl-β-D-galactopyranoside (MUG) substrate (Merck, Darmstadt, Germany) that was prepared in dimethyl sulfoxide (DMSO) and incubated for 60 min at room temperature in the dark. A reaction containing ABT buffer and the MUG substrate was used as negative control. After the incubation time, 20 µl of each reaction were transferred into the wells of a black 96-well plate containing 180 µl ABT buffer and fluorescence values were determined using Synergy^TM^ Mx microplate reader (BioTek) at 366 nm excitation and 445 nm emission wavelengths. A standard curve was obtained using 0.015625 µM to 4 µM of the fluorescent 4-methylumbelliferone (MU) standard. β-galactosidase units, or MUG units, were calculated as (pmol of substrate hydrolysed x dilution factor)/(culture volume in ml x OD_600_ x reaction time in minutes). The amount of hydrolysed substrate was determined from the standard curve as (emission reading – *y* intercept)/slope.

## Results

### Isolation of BAC-tolerant *L. monocytogenes* strains

The *L. monocytogenes* wildtype strain EGD-e was propagated on BHI agar plates supplemented with BAC to obtain genetically adapted strains. The wildtype strain could still grow in presence of 2 µg ml^-1^ BAC but was unable to grow on BHI agar plates containing higher BAC concentrations. However, single colonies appeared on plates containing 4 and 6 µg ml^-1^ BAC after 24 h, which likely acquired mutations to cope with the BAC stress. Since previous studies revealed that BAC-adapted *L. monocytogenes* isolates frequently mutate *fepR*, encoding the transcriptional regulator FepR, we first amplified the *fepR* gene and the *fepR* promoter region and analysed the sequence using Sanger sequencing. Indeed, all adapted strains acquired mutations in *fepR* or its promoter region (Fig. 1). The transcriptional regulator FepR possesses a helix-turn-helix (HTH) domain between residues 23 and 42, which is required for DNA binding. In addition, a putative substrate binding pocket was predicted to be located in the vicinity of residues 60, 100, 101, 104, 105, 119, 123, 126, 156, 159, 160 and 163 (Douarre *et al*. 2022). We identified seven BAC-tolerant strains with point mutations, amino acid insertions or deletions in the DNA binding site (S23L, L24F, INS29DIA, Δ45-46) and two with mutations or amino acid deletions in the putative substrate binding site (V115D, Δ99). Nine BAC-tolerant strains had mutations leading to a frameshift or the production of a truncated FepR protein (M126fs, W137fs, N170fs, Q140*, Y155*, G157*). We additionally isolated two suppressors that had base exchanges is the promoter region of the *fepRA* operon (G-27T and A-33G). All these mutations likely result in a reduced binding activity of the regulator or a decreased or abolished activity of FepR. For further analysis, we focussed on the BAC-tolerant strains EGD-e *fepR^Q140*^*, EGD-e *fepR^V115D^* and EGD-e *fepR^L24F^*. The TetR-family transcriptional regulator FepR represses the expression of the MATE family efflux pump FepA (Guérin *et al*. 2014). Hence, loss of function of the regulator subsequently leads to enhanced *fepA* expression. To verify that overproduction of FepA results in increased tolerance towards BAC, the IPTG-inducible plasmid pIMK3-*fepA* was constructed and introduced into the wildtype strain (*fepR^+^*). In addition, a *fepA* deletion strain (Δ*fepA*) was constructed to determine its tolerance towards BAC. Indeed, drop dilution assays revealed, that while the deletion of *fepA* led to slightly increased susceptibility towards BAC, overexpression of the transporter led to a significant increase in BAC tolerance, similar to that of the three selected *fepR* mutant strains (Fig. 2).

**Figure 1:**
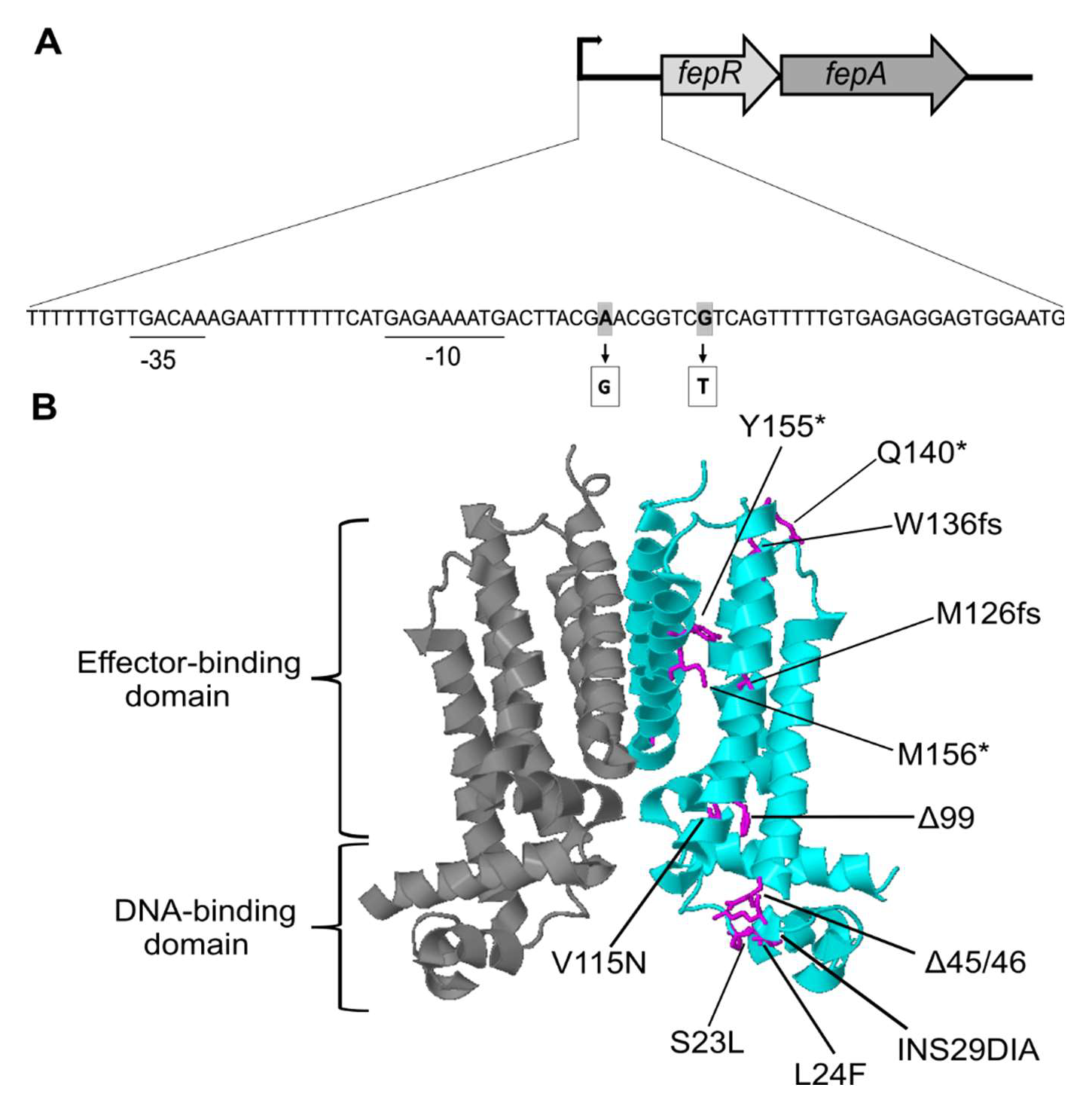
Mutations in the transcriptional regulator encoding gene *fepR*. **A** Genetic organization of the *fepRA* operon in *L. monocytogenes* EGD-e. The *fepRA* operon is composed of genes coding for the transcriptional regulator FepR and the MATE efflux pump FepA. The predicted promoter region is displayed along with the −10 and −35 regions (underlined). The base exchanges of the two suppressors in the *fepR* promoter are shown in grey. **B** The dimeric protein structure of the transcriptional regulator FepR (PDB: 5ZTC) was modified using Geneious Prime® v.2021.0.1. Single monomers are depicted in dark grey and cyan. Mutated amino acids are shown in magenta. Mutations leading to a stop codon are indicated with an asterisk and frameshift mutations are abbreviated with an fs. Amino acid insertions are indicated by INS and deletion by the Δ symbol.

**Figure 2:**
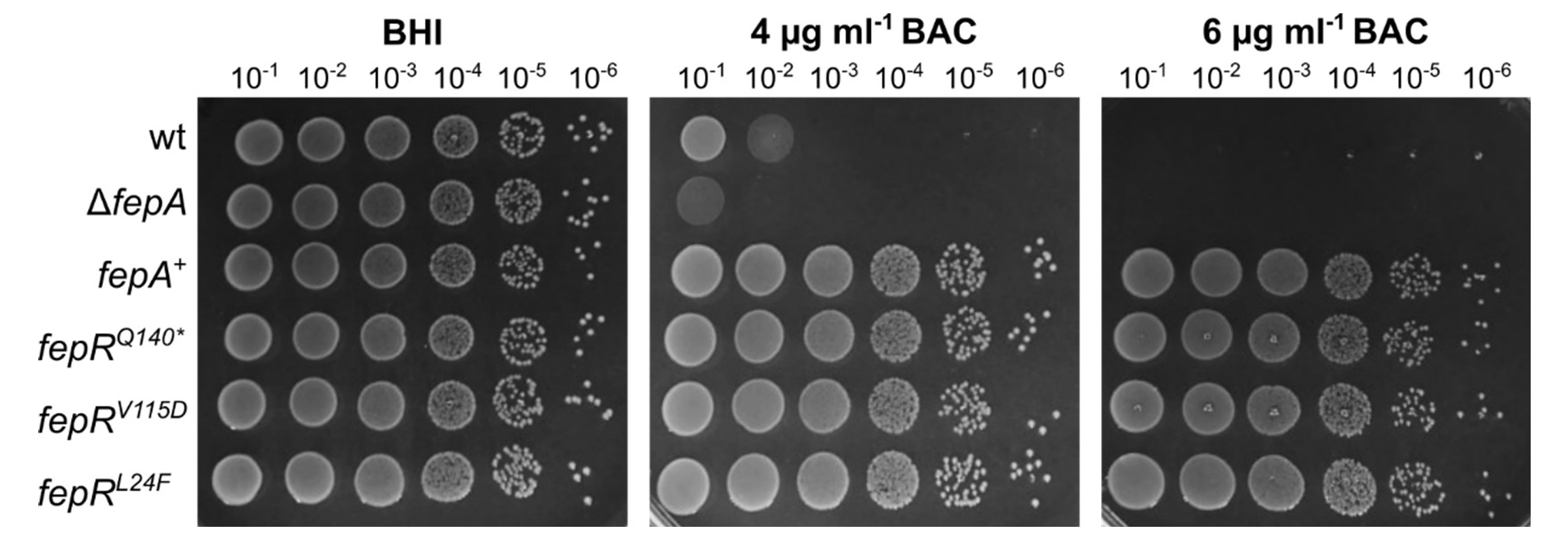
Increased BAC-tolerance of *fepR* suppressors. Drop dilution assays of *L. monocytogenes* strains EGD-e (wt), the *fepA* deletion strain LJR261 (Δ*fepA*), a wt strain containing the IPTG-inducible pIMK3-*fepA* plasmid LJR265 (*fepA*^+^), and the suppressor mutants LJR208 (*fepR^Q140*^*), LJR211 (*fepR^V115D^*) and LJR218 (*fepR^L24F^*). Cells were propagated on BHI plates or BHI plates containing 4 and 6 µg ml^-1^ BAC and plates incubated overnight at 37°C. Plates were supplemented with 1 mM IPTG to induce expression of FepA in the *fepA*^+^ strain. A representative image of at least three biological replicates is shown.

### Mutations in FepR alter DNA binding

According to structure predictions, the HTH motif of FepR, which is required for the interaction with DNA, is located between residues 23 and 42. Hence, we assumed that the mutation L24F has a negative effect on DNA binding by FepR. EMSA assays were performed to assess if DNA binding of FepR^L24F^ to the promoter region of the *fepRA* operon is altered. Indeed, in the concentration range used, no DNA-protein complexes could be detected for FepR^L24F^ in comparison to FepR^wt^ (Fig. 3A), indicating that the mutation L24F decreased DNA binding affinity of FepR. Binding of the two proteins to an unspecific DNA sequence from within the operon was not observed (Fig. 3A). We then tested the binding affinity of FepR^wt^ to the mutated promoter regions of the *fepRA* operon of the BAC-tolerant strains EGD-e *P_*fepR*_^G-27T^* and EGD-e *P_*fepR*_^A-33G^*. The base exchange from G to T 27 bp upstream of the start codon completely abolished binding of FepR^wt^ at the tested protein concentrations (Fig. 3B). Likewise, decreased binding affinity of FepR^wt^ was observed, when the promoter region contained an A to G substitution 33 bp before the start codon, but to a lesser extend then the G-27T promoter mutation (Fig. 3C). We further compared the promoter activity of the wildtype and mutated *fepR* promoters and determined their response to subinhibitory concentrations of BAC using β-galactosidase activity assays. Both *P_fepR_^G-27T^* and *P_fepR_*^A-33G^ showed significantly increased promoter activity in comparison to the wildtype promoter (Fig. 3D). *P_*fepR*_^G-27T^* hereby showed a higher activity than *P_fepR_*^A-33G^, which is in accordance with the difference in FepR binding capability (Fig. 3B-D). Increased β-galactosidase activity could be measured, when EGD-e *P_fepR_-lacZ* was grown in the presence of BAC (Fig. 3E), indicating that the expression of *fepRA* is slightly induced in response to BAC.

**Figure 3:**
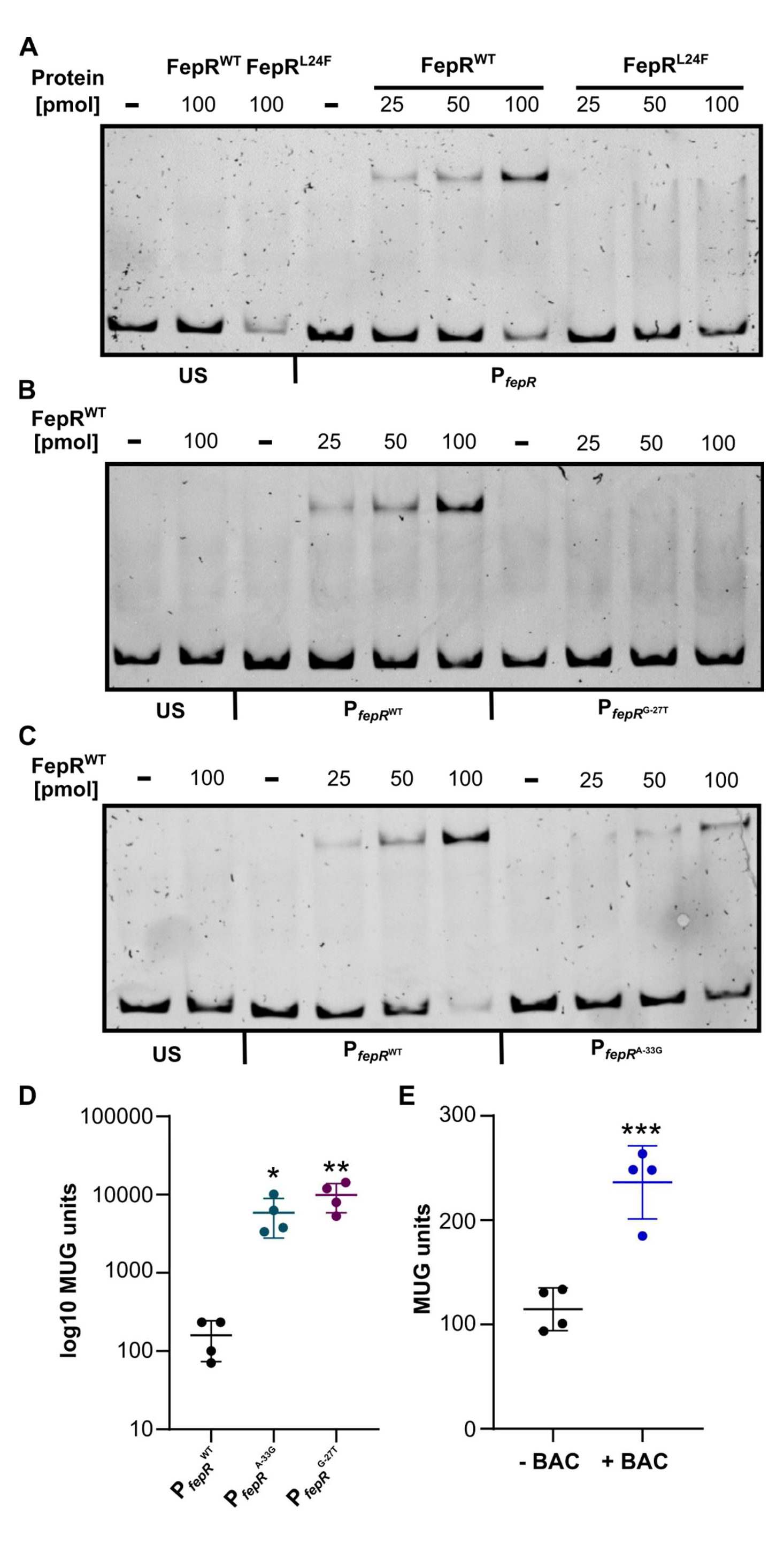
Interaction of FepR with the *fepRA* promoter and *P_fepR_* promoter activity. **A** Increasing concentrations of recombinant His-FepR^wt^ (lane 5-7) or His-FepR^L24F^ (lane 8-10) were incubated with a 150-bp fragment containing the promoter of the *fepRA* operon. **B** Increasing concentrations of FepR^wt^ were incubated with either the wildtype promoter region (*P_fepR_^wt^*) or the promoter region with a base exchange from G to T 27 bp upstream of the ATG (*P_fepR_^G-27T^*). **C** Incubation of FepR^wt^ with either *P_fepR_^wt^* or the promoter region with a base exchange from A to G 33 bp upstream of the ATG (*P_fepR_^A-33G^*). A short DNA sequence amplified from within the operon was incubated with 100 pmol FepR^wt^ or FepR^L24F^ and was included on each gel to exclude unspecific binding of the two proteins (US). Reactions without protein were used as an additional control (-). **D-E** Promoter activity assays. **D** EGD-e pPL3e-*P_fepR_^wt^*-*lacZ*, EGD-e pPL3e-*P_*fepR*_^A-33G^*-*lacZ* and EGD-e pPL3e-*P_*fepR*_^G-27T^*-*lacZ* were grown for 5 h in BHI medium and the promoter activity determined by β-galactosidase activity assays as described in the materials and methods section. Log10 of the MUG units are plotted to visualize the values obtained for EGD-e pPL3e-*P_fepR_^wt^*-*lacZ.* **E** Bacteria from a mid-logarithmic culture of strain EGD-e pPL3e- *P_fepR_^wt^*-*lacZ* were grown for 2 h in the presence or absence of 2.5 µg ml^-1^ BAC. The *P_fepR_* promoter activity was determined by β-galactosidase activity assays as described in the materials and methods section. For statistical analysis, one-way ANOVA coupled with Dunnett’s multiple comparison test was performed (* *p* ≤ 0.05; ** *p* ≤ 0.01; *** *p* ≤ 0.001).

### Ethidium bromide is a substrate for the efflux pump FepA

Previous work has shown that the deletion of *fepR* resulted in an increased ethidium bromide (EtBr) resistance (Guérin *et al*. 2014). To assess, whether this resistance can be explained by the overproduction of FepA, an EtBr accumulation assay was performed with the wildtype, Δ*fepA*, *fepA^+^*, the EGD-e *fepR^Q140*^* suppressor mutant and a Δ*fepA* complementation strain (CΔ*fepA*). This assay revealed that a strain lacking the efflux pump FepA accumulated more EtBr as the wildtype strain (Fig. 4). In contrast, the more gradual slope of the EGD-e *fepR^Q140*^* suppressor indicates slightly reduced accumulation. Similarly, a significantly reduced accumulation of EtBr was observed for the Δ*fepA* complementation strain and the *fepA* overexpression strain (*fepA^+^*) (Fig. 4), suggesting that FepA is able to export EtBr.

**Figure 4:**
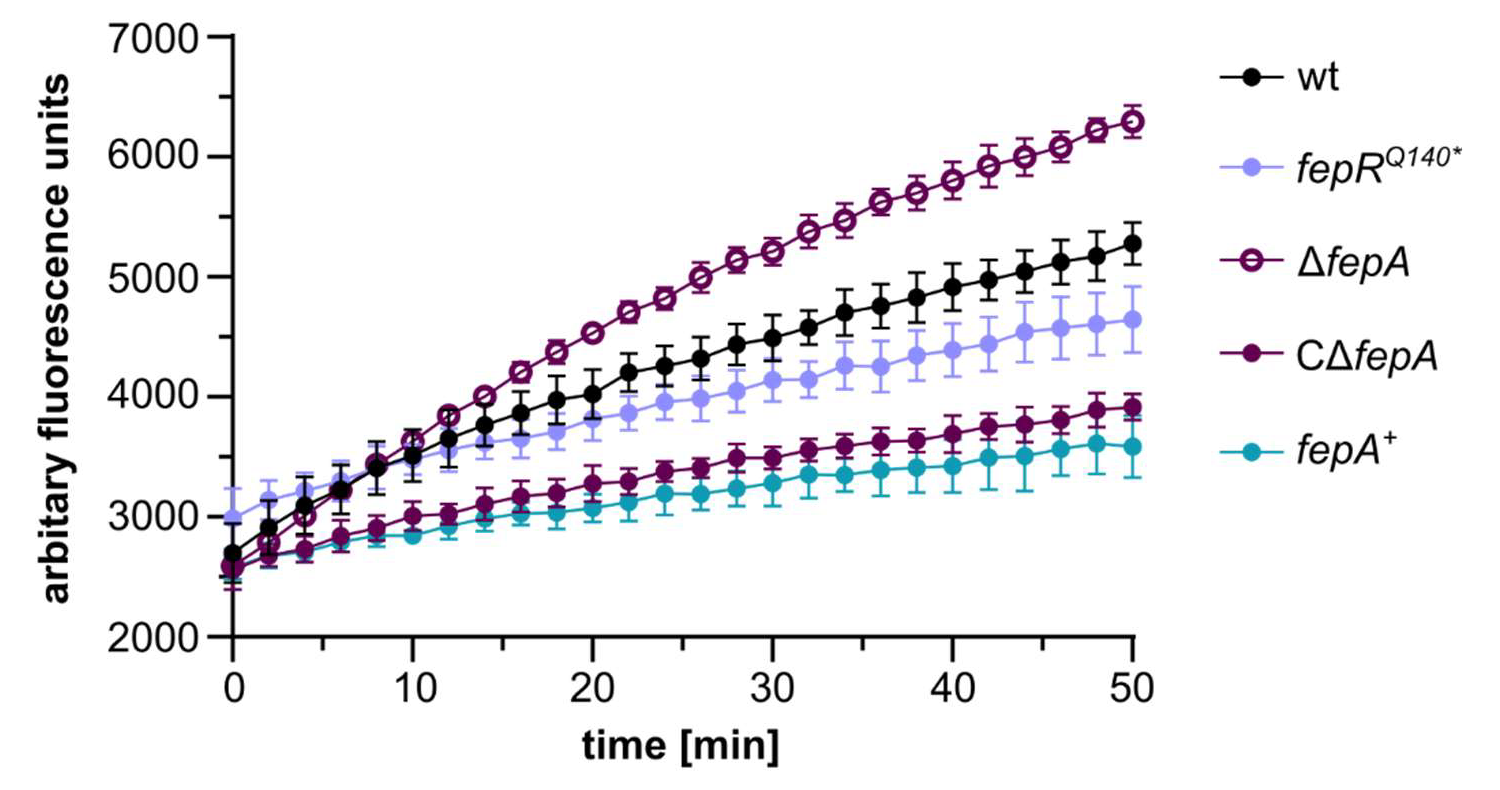
EtBr accumulation assay. Accumulation of ethidium bromide (EtBr) by *L. monocytogenes* EGD-e (wt), the *fepR^Q140*^* suppressor strain (LJR208), a Δ*fepA* mutant strain (LJR261), a Δ*fepA* complementation strain (CΔ*fepA*) and a *fepA* overexpression strain (*fepA*^+^) was measured at an excitation wavelength of 500 nm and an emission wavelength of 580 nm for 50 min. The average values and standard deviations of three independent experiments are depicted.

### FepA contributes to cross-resistance and tolerance towards CTAB

The MATE efflux pump FepA has been previously associated with fluoroquinolone resistance. We thus wondered whether the BAC-tolerant strains are also more tolerant towards other antimicrobials such as the fluoroquinolone antibiotic ciprofloxacin, the aminoglycoside gentamycin, the cephalosporin cefuroxime or the surfactant CTAB. Indeed, growth experiments revealed that the mutations in *fepR*, and hence overexpression of FepA, additionally conferred resistance towards ciprofloxacin, gentamycin and increased tolerance towards CTAB (Fig. 5), but not towards cefuroxime (data not shown). Similar results were obtained for the *fepA^+^* strain, which artificially overproduces FepA (Fig. 5). Interestingly, we observed suppressor formation in the wildtype strain on plates containing CTAB, which hasn’t been described so far.

**Figure 5:**
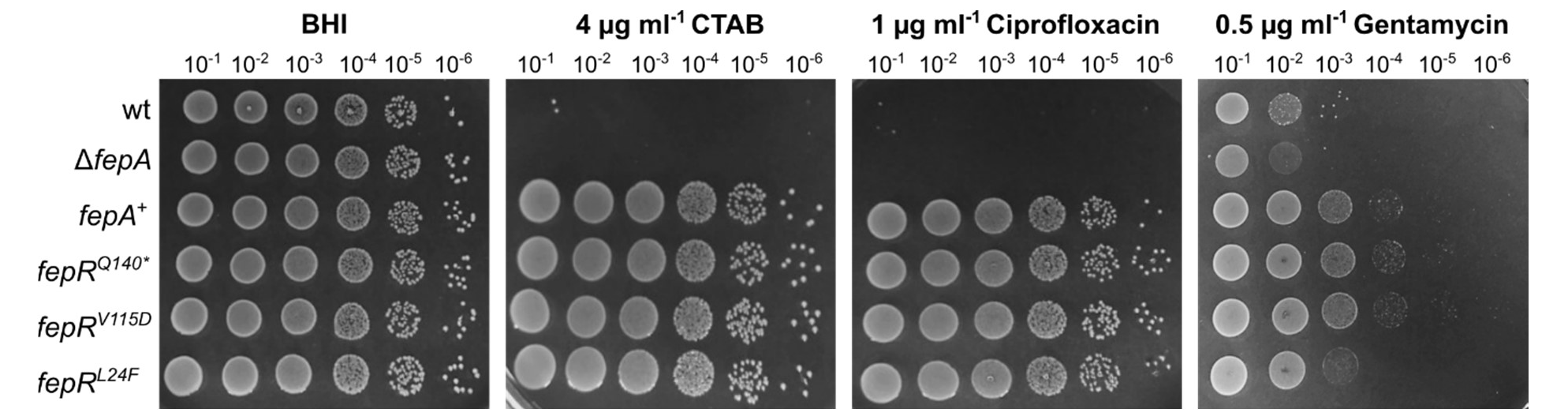
Cross-resistance of *fepR* mutant strains. Drop dilution assays of *L. monocytogenes* strains EGD-e (wt), the *fepA* deletion strain LJR261 (Δ*fepA*), a wt strain containing the IPTG-inducible pIMK3-*fepA* plasmid LJR265 (*fepA*^+^), and the suppressor mutants LJR208 (*fepR^Q140*^*), LJR211 (*fepR^V115D^*), and LJR218 (*fepR^L24F^*). Cells were propagated on BHI plates or BHI plates containing 4 µg ml^-1^ CTAB, 1 µg ml^-1^ ciprofloxacin or 0.5 µg ml^-1^ gentamycin. All plates were supplemented with 1 mM IPTG to induce expression of *fepA* in the *fepA*^+^ strain. Plates were incubated overnight at 37°C. A representative image of at least three biological replicates is shown.

### Mutations in *sugR* confer resistance towards CTAB

For the isolation of CTAB-tolerant *L. monocytogenes* strains, the wildtype strain EGD-e was propagated onto BHI plates containing varying CTAB concentrations. Growth of the wildtype was diminished at concentrations above 1 µg ml^-1^ CTAB and CTAB-tolerant strains were isolated in the presence of 2 and 4 µg ml^-1^ of CTAB. The genome sequence of 2 and 7 of the CTAB-tolerant strains isolated from 2 and 4 µg ml^-1^, respectively, were determined by whole genome sequencing to identify the underlying mutations. All of these strains carried mutations in the coding or promoter region of *sugR*, encoding a TetR-family transcriptional regulator (Fig. 6A-B). 4 of the CTAB-tolerant strains had a 1 bp deletion leading to a frameshift after phenylalanine at position 49, 4 strains carried point mutations leading to a premature stop after 64 or 122 amino acids and one strain carried a point mutation in the *sugR* promoter region. SugR is encoded in an operon together with *sugE1* and *sugE2*, coding for two SMR efflux pumps. Deletion of the regulator *sugR* leads to the overexpression of SugE1 and SugE2 and by this to an increased tolerance towards QACs, including BAC and CTAB (Jiang *et al*. 2020). However, to our knowledge, this is the first time that strains were isolated that acquired mutations in *sugR* under CTAB treatment. To assess the impact of SugE1 and SugE2 on BAC and CTAB tolerance, drop dilution assays were performed with a strain lacking both efflux pumps, a *sugE1*/*2* overexpression strain (*sugE1*/*2*^+^), the two CTAB-tolerant strains EGD-e *sugR^D122*^* and EGD-e *sugR^F49fs^*, as well as the wildtype strain. The BAC-tolerant strain EGD-e *fepR*^Q140*^ was included as a control. *sugE1*/*2*^+^, as well as the two CTAB-tolerant strains showed a growth advantage on BHI plates supplemented with CTAB and BAC in comparison to the wildtype and Δ*sugE1*/*2* deletion strains. However, EGD-e *sugR^D122*^* and EGD-e *sugR^F49fs^* were unstable in the presence of BAC as they readily formed suppressors, while the growth of EGD-e *fepR^Q140*^* was not affected by BAC (Fig. 6C). Our results therefore indicate that FepA and SugE1/2 are the dominant efflux pumps for BAC and CTAB, respectively. In comparison to the *fepR-* associated suppressors, no cross-resistance towards gentamycin and ciprofloxacin was observed in association with the overproduction of SugE1/2 (Fig. S1).

**Figure 6:**
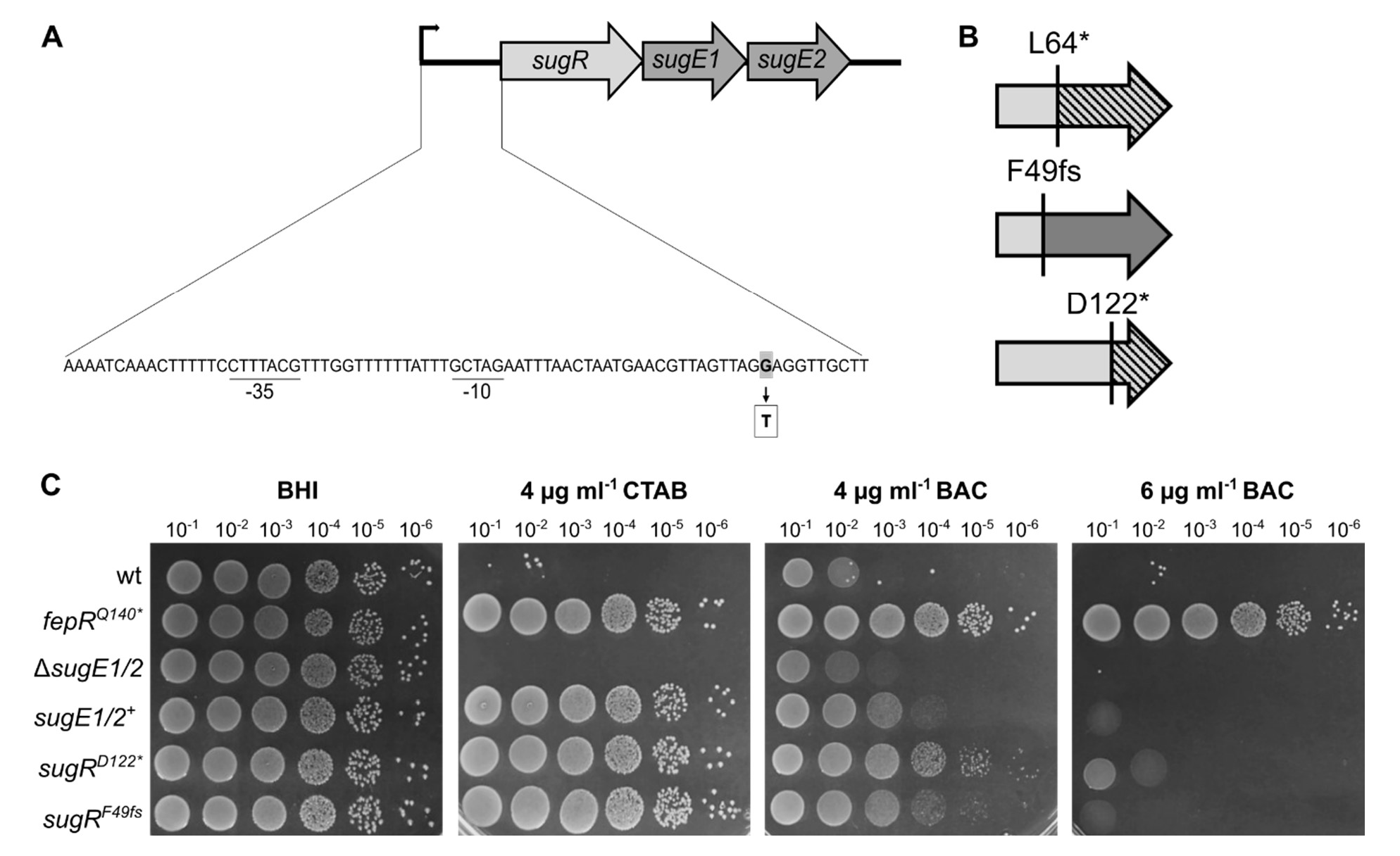
Acquired CTAB tolerance due to mutations in *sugR*. **A** Genetic organization of the *sug* operon in *L. monocytogenes* EGD-e with the predicted promoter region including the −10 and −35 regions (underlined). The *sug* operon contains genes coding for the transcriptional regulator SugR and the two SMR efflux pumps SugE1 and SugE2. Base exchange from one of the suppressors is displayed in grey (adapted from Jiang *et al*., 2020). **B** Mutations in the *sugR* gene (depicted in light grey) in strains isolated in presence of CTAB. Dark grey colour depicts part of the gene/protein that is affected by the frameshift. The deleted parts of the protein are shown as dashed lines. **C** Drop dilution assays of *L. monocytogenes* strains EGD-e (wt), a *sugE1/2* deletion strain (Δ*sugE1/2*), a wt strain containing the IPTG-inducible pIMK3-*sugE1/2* plasmid LJR301 (*sugE1/2*^+^), and the suppressor mutants *sugR^D122*^*(LJR248), and *sugR^F49fs*^* (LJR258). The *fepR^Q140*^* suppressor mutant (LJR208) was used as a control. Cells were propagated on BHI plates or BHI plates containing 4 µg ml^-1^ CTAB, 4 and 6 µg ml^-1^ BAC. All plates were supplemented with 1 mM IPTG to induce the expression of *sugE1/2* in the *sugE1/E2*^+^ strain. Plates were incubated overnight at 37°C. A representative image of at least three biological replicates is shown.

### SugE1/2 and FepA can partially compensate for each other in presence of biocide stress

In an attempt to identify further tolerance mechanisms, the two deletion strains Δ*fepA* and Δ*sugE1/2* were again adapted to BAC and CTAB. Δ*sugE1/2* readily formed suppressors in presence of 6 µg ml^-1^ BAC and 4 µg ml^-1^ CTAB within a day. The Δ*fepA* strain evolved suppressors in presence of 4 and 6 µg ml^-1^ CTAB within a day, while it took two days to isolate BAC-tolerant suppressors (5 µg ml^-1^ BAC). To assess if overproduction of FepA and SugE1/2 can compensate for the lack of SugE1/2 or FepA, respectively, isolated suppressors were screened for mutations in the respective transcriptional regulator. Indeed, all isolated Δ*sugE1/2* suppressors carried mutations in the *fepR* gene, regardless of the selective pressure (Fig. 7C). Similarly, all Δ*fepA* isolates had mutations in the *sugR* gene, most of which led to either a premature stop or a frameshift. A similar growth behaviour could be observed for the Δ*fepA* and Δ*sugE1/2* suppressors as compared to the BAC- and CTAB-tolerant wildtype strains (Figs. 2, 5 and 7A-B). Δ*fepA sugR^L57*^* and Δ*fepA sugR^F49fs^* showed enhanced tolerance in presence of 4 µg ml^-1^ CTAB and 4 µg ml^-1^ BAC, while only minor growth was observed in presence of 6 µg ml^-1^ BAC.

**Figure 7:**
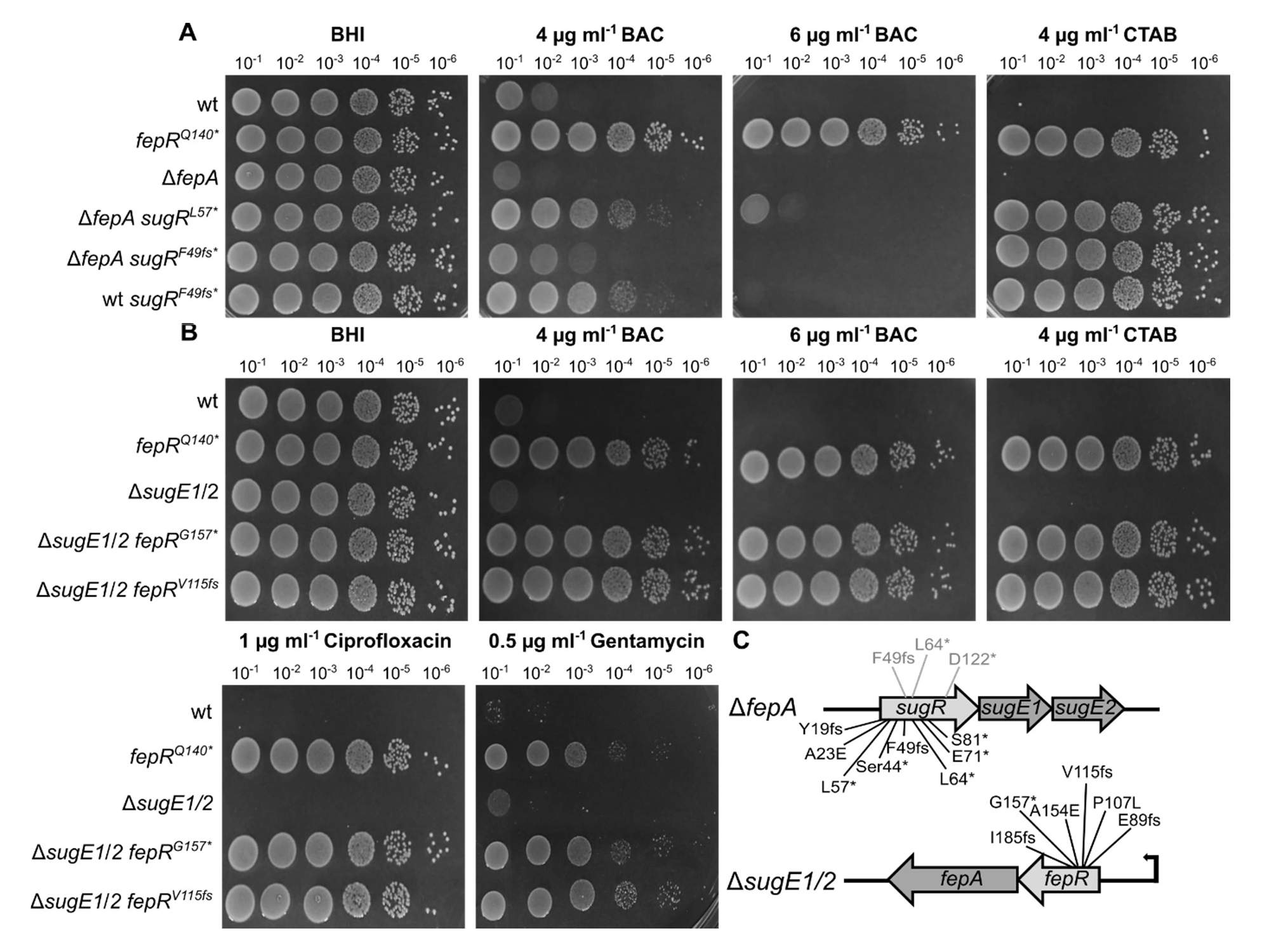
SugE1/2 and FepA can partially compensate for each other in the presence of biocide stress. **A** Drop dilution assays of *L. monocytogenes* strains EGD-e (wt), the *fepA* deletion strain (Δ*fepA*) and the suppressor mutants Δ*fepA sugR^L57^*\*(LJR280), Δ*fepA sugR^F49fs^* (LJR267) and wt *sugR^F49fs^* (LJR258). The *fepR^Q140*^* suppressor mutant (LJR208) was used as a control. Cells were propagated on BHI plates or BHI plates containing 4 and 6 µg ml^-1^ BAC or 4 µg ml^-1^ CTAB and incubated overnight at 37°C. **B** Drop dilution assays of *L. monocytogenes* strains EGD-e (wt), the *sugE1/2* deletion strain (Δ*sugE1*/*2*), and the suppressor mutants Δ*sugE1*/*2 fepR^G157*^* (LJR270), and Δ*sugE1*/*2 fepR^V115fs^*(LJR276), isolated on CTAB and BAC, respectively. The *fepR^Q140*^* suppressor mutant was used as a control. Cells were propagated on BHI plates or BHI plates supplemented with 4 and 6 µg ml^-1^ BAC, 4 µg ml^-1^ CTAB, 1 µg ml^-1^ ciprofloxacin or 0.5 µg ml^-1^ gentamycin and incubated overnight at 37°C. A representative image of at least three biological replicates is shown. **C** Acquired *sugR* and *fepR* mutations in the Δ*fepA* or Δ*sugE1*/*2* background, respectively. The mutations that were previously identified in the wildtype background are depicted in grey.

Interestingly, the wildtype strain harbouring the F49fs mutation in *sugR* seems to be slightly more tolerant towards BAC than the corresponding Δ*fepA* strain (Fig. 7A). In addition, no cross-resistance towards ciprofloxacin and gentamycin was observed for Δ*fepA sugR^L57*^* and Δ*fepA sugR^F49fs^* (Fig. S2). In contrast, Δ*sugE1/2 fepR^G157*^* and Δ*sugE1/2 fepR^V115fs^* showed similar growth as the wildtype strain carrying the *fepR^Q140*^* mutation, where not only increased tolerance towards BAC and CTAB was observed, but also cross-resistance towards ciprofloxacin and gentamycin (Fig. 7B). These results indicate that FepA and SugE1/2 can at least partially compensate for each other.

### Strains lacking the two major QAC efflux systems can still acquire tolerance

To assess whether *L. monocytogenes* possesses a third mechanism to adapt to QACs, the Δ*fepA* Δ*sugE1/2* double deletion strain was constructed and propagated in the presence of BAC and CTAB. Interestingly, no suppressor formation was observed on BHI plates containing 6 µg ml^-1^ BAC or 4 µg ml^-1^ CTAB, which were previously used to isolate BAC- und CTAB-tolerant strains, likely due to the absence of two important efflux systems. However, the Δ*fepA*Δ*sugE1/2* deletion strain could still adapt to 5 µg ml^-1^ BAC and 2 µg ml^-1^ CTAB. Genomic alterations for two BAC- and two CTAB-tolerant strains were determined by whole genome sequencing. Interestingly, all tolerant isolates acquired mutations in *lmo1753* encoding a putative diacylglycerol kinase and Sanger sequencing of additional mutants likewise identified mutations in *lmo1753*. The phenotype of the suppressors slightly varied in presence of different stresses. The two CTAB-tolerant strains Δ*fepA*Δ*sugE1/2* CTAB1 and Δ*fepA*Δ*sugE1/2* CTAB2 showed only slightly increased tolerance on plates containing 2 µg ml^-1^ CTAB and 4 µg ml^-1^ BAC. In contrast, the BAC-tolerant strain Δ*fepA*Δ*sugE1/2* BAC2 could grow to some extent in the presence of CTAB and showed enhanced growth even in the presence of up to 6 µg ml^-1^ BAC as compared to the wildtype strain and the Δ*fepA*Δ*sugE1/2* deletion strain (Fig. 8). Apart from the mutation in *lmo1753*, this suppressor carried a mutation in the promoter region of *lmo1682*, encoding a putative multidrug efflux pump, whose overexpression is likely responsible for the increased BAC tolerance. The BAC-tolerant strain Δ*fepA*Δ*sugE1/2* BAC1 also showed an enhanced tolerance towards BAC as compared to the Δ*fepA*Δ*sugE1/2* deletion strain, however, no growth advantage could be observed on BHI plates containing CTAB (Fig. 8). Interestingly, Δ*fepA*Δ*sugE1/2* BAC1 was more sensitive towards the antibiotic cefuroxime in comparison to the parental and wildtype strain as well as an increased tolerance to gentamycin similar to EGD-e *fepR^Q140*^* (Fig. 8). Remarkably, we could neither identify any additional mutations for Δ*fepA*Δ*sugE1/2* BAC1 apart from the mutation in *lmo1753*, which is identical to the mutation in Δ*fepA* Δ*sugE1/2* CTAB1 and CTAB2, nor indications of gene amplification events that could explain the distinct phenotype. No cross-adaption towards ciprofloxacin was observed for any of the Δ*fepA*Δ*sugE1/2* suppressors (Fig. 8).

**Figure 8:**
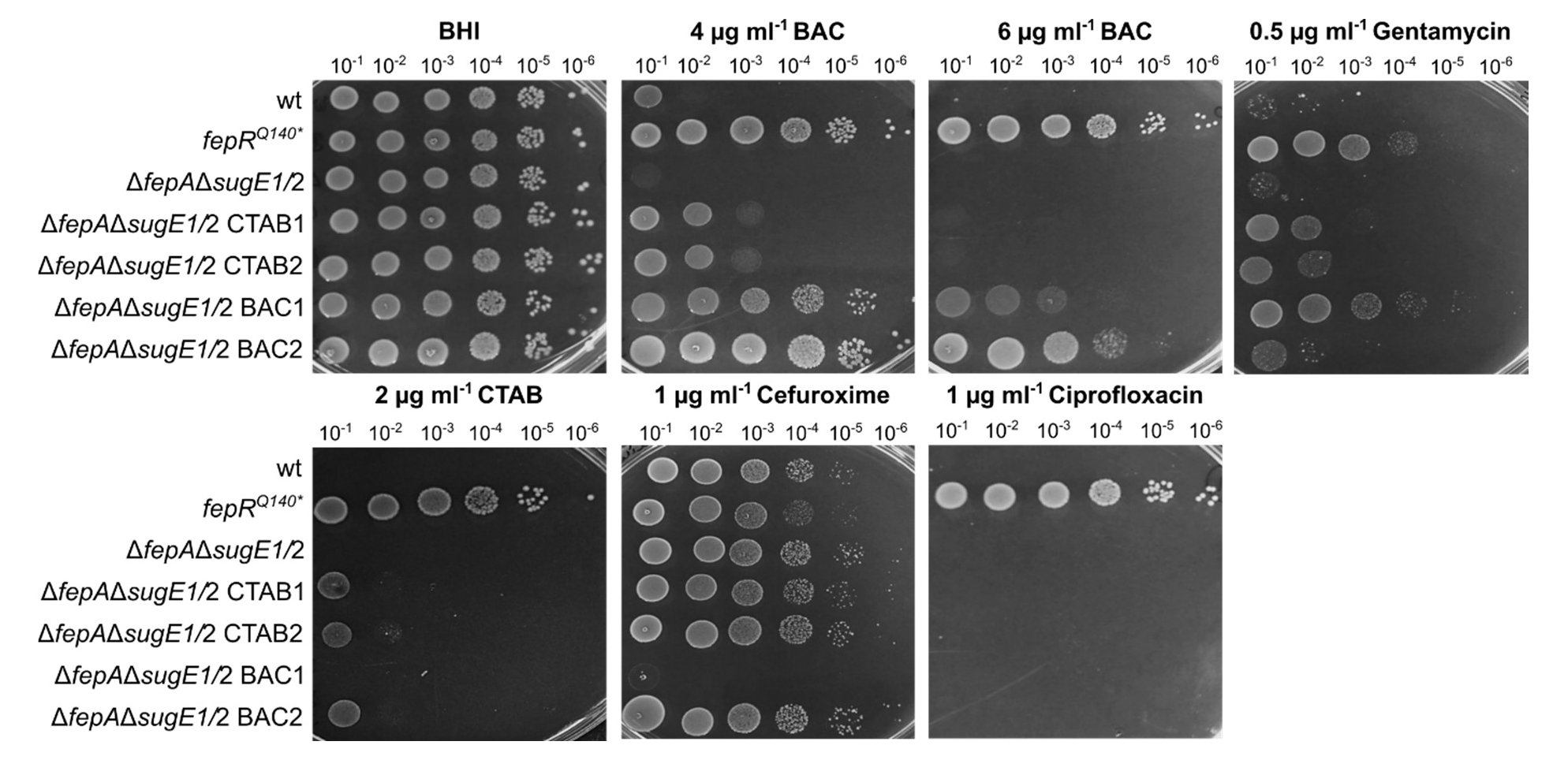
QAC tolerance of the Δ*fepAΔsugE1/2* deletion strain. Drop dilution assays of *L. monocytogenes* strains EGD-e (wt), the *fepA sugE1/2* deletion strain (Δ*fepA*Δ*sugE1/2*), the two suppressor mutants that were isolated in presence of CTAB, LJR327 (Δ*fepA*Δ*sugE1/2 lmo1753^K19fs^,* short CTAB1) and LJR328 (Δ*fepA*Δ*sugE1/2 lmo1753^K19fs^,* short CTAB2), and the two suppressor mutants that were isolated in presence of BAC, LJR326 (Δ*fepA*Δ*sugE1/2 lmo1753^K19fs^,* short BAC1) and LJR330 (Δ*fepA*Δ*sugE1/2 lmo1753^V225fs^ P_*lmo1682*_^G-37A^*, short BAC2). A representative image of at least three biological replicates is shown.

## Discussion

Survival and proliferation of *L. monocytogenes* in the food industry is an ongoing concern, and while there are various countermeasures to combat the contamination of food products, such as osmotic stress, extreme temperatures or the use of disinfectants, the pathogen still successfully manages to withstand the harsh conditions present in food-processing facilities, resulting in reoccurring outbreaks. To counteract the spread of *L. monocytogenes,* it is crucial to understand and elucidate the underlying mechanism that permit their successful evasion. Outbreaks are often associated with strains that tolerate below working concentrations of QACs, such as BAC or CTAB, the most commonly used active agents in disinfectants (Weber *et al*. 2007). In this study, we assessed the ability of the laboratory wildtype strain EGD-e to adapt to low levels of BAC and CTAB under laboratory growth conditions. Since previous studies have focused on the analyses of *L. monocytogenes* isolates, which exhibit a high frequency of genomic variations, our findings using the laboratory model strain represent a more generalized assessment. While previously isolated strains often merely acquired a transient tolerance towards QACs that was lost after passaging of the strains in absence of the stress, our strains readily formed stable suppressors that allowed growth in presence of BAC and CTAB.

In our study, BAC-tolerant *L. monocytogenes* strains exclusively carried mutations in *fepR*, which encodes a transcriptional regulator. These findings are in accordance with previous studies, which focussed on serial adapted isolated *L. monocytogenes* and *Listeria spp.* strains that were exposed to BAC or ciprofloxacin stress and likewise found that the majority of strains acquired mutations in *fepR* (Bland *et al*. 2022; Bolten *et al*. 2022; Douarre *et al*. 2022). FepR is a TetR-like transcriptional regulator that negatively regulates the *fepRA* operon, which encodes the regulator itself as well as the MATE family efflux pump FepA (Guérin *et al*. 2014). Strains that carried a mutation in *fepR*, as well as strains that artificially overexpress the efflux pump FepA exhibited increased tolerance not only towards BAC, but also towards CTAB, ciprofloxacin and gentamycin, indicating that extrusion by the transporter is rather unspecific. This observation further supports the previous hypothesis that de-repression of *fepA* is the reason for the observed QAC tolerance (Guérin *et al*. 2014; Bland *et al*. 2022; Bolten *et al*. 2022; Douarre *et al*. 2022). We further substantiated this hypothesis by showing that a mutation in the DNA binding domain of FepR resulted in decreased binding to the promoter of the *fepRA* operon in comparison to the wildtype FepR protein. Likewise, mutations in the *fepRA* promoter region resulted in reduced binding of the wildtype FepR and thus, to increased promoter activity. This enhanced promoter activity could then result in an enhance production of FepA and subsequent export of BAC. Interestingly, we did not identify any mutations in either of the two chromosomally located efflux pumps MdrL or Lde that were previously described to be involved in QAC adaptation and whose expression is commonly upregulated in tolerant *L. monocytogenes* isolates (Mata *et al*. 2000; Jiang *et al*. 2019; Jiang *et al*. 2012; Godreuil *et al*. 2003). This was the case even in the absence of *fepA* and/or *sugE1/2*, suggesting that neither play a significant role in BAC or CTAB tolerance under the tested conditions. Altogether, we can conclude that the acquisition of mutations in *fepR* and the associated elevation of FepA levels and activity is the dominant mode of tolerance towards BAC in the EGD-e wildtype strain. In contrast, suppressors isolated in presence of CTAB stress solely acquired mutations in the *sugR* gene, coding for a different transcriptional regulator. Unexpectedly, none of the isolated suppressors carried mutations in the *fepR* gene. SugR is involved in the repression of the two SMR efflux pumps SugE1 and SugE2 that were previously shown to confer tolerance towards QACs such as BAC, CTAB or didecyldimethylammonium chloride (DDAC) in *L. monocytogenes*. Accordingly, expression of the efflux system was shown to be induced in presence of BAC, suggesting that BAC can inhibit the SugR-dependent repression of *sugE1/2* (Jiang *et al*. 2020). Similarly to previous findings, the overexpression of SugE1/2 either due to mutations in its repressor or artificially induced did not result in any further cross-adaption in contrast to suppressors with *fepR* mutations (Guérin *et al*. 2014; Bland *et al*. 2022; Jiang *et al*. 2020). Cross-resistance was also observed for *Pseudomonas aeruginosa* and *E. coli* isolates after BAC adaptation. While *P. aeruginosa* isolates acquired tolerance towards polymyxin B and other, BAC-adapted *E. coli* strains exhibited increased MIC for ampicillin and/or ciprofloxacin (Kim *et al*. 2018; Nordholt *et al*. 2021). This raises the question, if CTAB should be used more frequently in commercial disinfectants than BAC to prevent the emergence of multi-resistant strains. We also found that although both efflux systems could compensate for the loss of the other, overexpression of FepA results in a higher BAC-tolerance than overexpression of SugE1/2, suggesting that the two efflux systems do not have specific substrates, but that the affinity seems to differ for the two QACs, BAC and CTAB. We further evolved a strain that lacks the two major QAC efflux systems FepA and SugE1/2 to identify additional tolerance mechanisms. To our surprise, all isolated suppressors acquired mutations in *lmo1753,* which does not code for an additional efflux system. Instead, *lmo1753* shares 64% sequence identity and 87.2% similarity with the gene coding for the diacylglycerol kinase DgkB from *Bacillus subtilis*, which contributes to the biosynthesis of lipoteichoic acids (LTA) by recycling the toxic intermediate phosphatidic acid (Jerga *et al*. 2007; Matsuoka *et al*. 2011). This finding indicates that SugE1/2 and FepA are the key BAC and CTAB efflux systems in the *L. monocytogenes* wildtype strain EGD-e. LTAs make up a great portion of the Gram-positive cell wall and have been shown to play crucial roles in cellular growth, morphology and division. They are anchored to the cell membrane and mainly consist of a polyglycerolphosphate backbone that contributes to the overall negative surface charge of the cell (Campeotto *et al*. 2014). The negatively charged backbone can be masked by decoration with positively charged D-alanylation. This decoration can be rather flexible and can fluctuate according to environmental and cellular cues, allowing adjustment of the cellular surface charge and hence variation in the cation homeostasis of the membrane (Percy and Gründling 2014). A decrease in cellular surface charge was previously associated with the survival of adapted *E. coli* strains in the presence of BAC, as they carried mutations in *lpxM*, encoding an enzyme involved in lipid A biosynthesis (Nordholt *et al*. 2021). Likewise, an increased negative surface charge was shown to be beneficial in a high-level BAC-tolerant *Pseudomonas fluorescence* strain (Nagai *et al*. 2003). It is tempting to speculate that mutations in *lmo1753* result in altered LTA synthesis followed by a distorted negative surface charge subsequently hindering binding of the positively charged head groups of BAC and CTAB. While the activity of DgkB has often mainly been discussed in the context of LTA biosynthesis, phosphatidic acid can also be utilized for the production of other glycolipids and phospholipids, including cardiolipin, lysyl-phosphatidylglycerol or phosphatidylethanolamine. Likewise, diacylglycerol, the substrate of DgkB, is aside from LTA biosynthesis also crucial for the production of triglucosyldiacyl-glycerol (Hashimoto *et al*. 2013). Hence, aberrant DgkB activity might generally result in an altered lipid profile. Besides efflux systems, changes in fatty acid composition and concomitant altered membrane fluidity have been proposed to contribute to QAC tolerance in several organisms. A study from 2002 described a tolerant *L. monocytogenes* isolate that showed a slight shift in the length of fatty acids (To *et al*. 2002). General alterations of the fatty acid profile and content was likewise associated with QAC tolerance in *Serratio marcescens* (Chaplin 1952) and *P. aeruginosa* (Guerin-Mechin *et al*. 2000; Jones *et al*. 1989). It remains elusive how mutations in *lmo1753* contribute to QAC tolerance, but our findings highlight the ability of *L. monocytogenes* to adapt to QAC *via* an export-independent mechanism. One of the Δ*fepA* Δ*sugE1/2* suppressors showed enhanced tolerance towards BAC in comparison to the other isolated mutants with the same genetic background. Interestingly, the strain acquired in addition to the mutation in *lmo1753* a mutation in the promoter region of *lmo1683,* which encodes a putative major facilitator family transporter. To our knowledge, no function was assigned for this transporter so far; however, our study suggests that it might be involved in the export of BAC. Further analysis of the transporter is required to elucidate its role in the efflux of QACs.

It has to be mentioned, that all suppressors were isolated on BHI complex medium and in the presence of below working concentrations of BAC and CTAB. Those rather ideal conditions are not commonly found in food-processing facilities. However, similar BAC concentrations are often found in hard to reach places, when disinfectants are not properly applied and concentrations of approximately 0.5 µg ml^-1^ were for instance reported in household wastewater, creating an environment that allows adaptation of the pathogen prior to entering food-processing plants (Tezel and Pavlostathis 2015).

Altogether, our study supported previous findings that designated the efflux pump FepA as the major BAC extrusion system in *L. monocytogenes*. We further showed that SugE1/2 play a similar role for CTAB-tolerance and that both systems are the two main efflux systems for BAC and CTAB. Our suppressor screen also revealed the ability of *L. monocytogenes* to acquire tolerance independent of the presence and/or overexpression of efflux systems, likely due to alterations in the lipid profile, which will be further analysed in the future.

## Acknowledgements

We are grateful to Julia Busse for technical assistance and thank Prof. Jörg Stülke for providing laboratory space, equipment and consumables and to the Göttingen Center for Molecular Biosciences (GZMB) for financial support.

## Author Contributions

LMS: Investigation, formal analysis, conceptualization, methodology, supervision, validation, writing – original draft. FD: Investigation, formal analysis, writing – review & editing. LMdSM: Investigation, formal analysis, writing – review & editing. JR: Investigation, formal analysis, conceptualization, supervision, funding acquisition, writing – original draft.

