## Supplementary Material for "Adaptation mechanisms of *Listeria monocytogenes* to quaternary ammonium compounds"

**Running title:** Adaptation of *L. monocytogenes* to biocides

**Supplementary Tables**

**Table S1: Bacterial strains used in this study**

| **Unique ID** | **Strain name and resistance** | **Source** |
|---|---|---|
| ***Escherichia coli* strains** | | |
| ANG124 | DH5α pKSV7; AmpR | (Smith and Youngman 1992) |
| ANG4243 | XL1-Blue pIMK3; KanR | (Monk *et al.* 2008) |
| ANG5181 | XL1-Blue pPL3e-*lacZ*; CamR | (Rismondo *et al.* 2019) |
| EJR149 | XL10-Gold pWH844; AmpR | (Schirmer *et al.* 1997) |
| EJR227 | XL10-Gold pIMK3-*fepA*; KanR | This study |
| EJR229 | XL1-Blue pKSV7-Δ*sugE1/2*; AmpR | This study |
| EJR230 | XL1-Blue pKSV7-Δ*fepA*; AmpR | This study |
| EJR242 | XL10-Gold pWH844-*fepR*; AmpR | This study |
| EJR248 | XL10-Gold pWH844-$fepR^{L24F}$; AmpR | This study |
| EJR257 | DH5α pPL3e-$P_{fepR}$-*lacZ*; CamR | This study |
| EJR258 | DH5α pPL3e-$P_{fepR}^{A-33G}$-*lacZ*; CamR | This study |
| EJR259 | XL10-Gold pIMK3-*sugE1/2*; KanR | This study |
| EJR260 | DH5α pPL3e-$P_{fepR}^{G-27T}$-*lacZ*; CamR | This study |
| ***Listeria monocytogenes* strains** | | |
| ANG873 | EGD-e | (Glaser *et al.* 2001) |
| LJR187 | EGD-e $fepR^{N170fs}$ | This study |
| LJR188 | EGD-e $P_{fepR}^{G-27T}$ | This study |
| LJR190 | EGD-e $fepR^{G157*}$ | This study |
| LJR194 | EGD-e $fepR^{\Delta 45-46}$ | This study |
| LJR196 | EGD-e $fepR^{\Delta 99}$ | This study |
| LJR208 | EGD-e $fepR^{Q140*}$ | This study |
| LJR209 | EGD-e $fepR^{Y155*}$ | This study |

| | | |
|---|---|---|
| LJR210 | EGD-e *fepR*$^{Q140*}$ | This study |
| LJR211 | EGD-e *fepR*$^{V115D}$ | This study |
| LJR212 | EGD-e *fepR*$^{\Delta 45\text{-}46}$ | This study |
| LJR213 | EGD-e *fepR*$^{S23L}$ | This study |
| LJR214 | EGD-e *fepR*$^{Q140*}$ | This study |
| LJR215 | EGD-e $P_{fepR}{}^{A\text{-}33G}$ | This study |
| LJR216 | EGD-e *fepR*$^{M126fs}$ | This study |
| LJR217 | EGD-e *fepR*$^{\Delta 45\text{-}46}$ | This study |
| LJR218 | EGD-e *fepR*$^{L24F}$ | This study |
| LJR219 | EGD-e *fepR*$^{INS29DIA}$ | This study |
| LJR220 | EGD-e *fepR*$^{\Delta 45\text{-}46}$ | This study |
| LJR221 | EGD-e *fepR*$^{W137fs}$ | This study |
| LJR222 | EGD-e *fepR*$^{M126fs}$ | This study |
| LJR231 | EGD-e pIMK3-*fepA*; KanR | This study |
| LJR234 | EGD-e $P_{sugR}{}^{G\text{-}11T}$ | This study |
| LJR235 | EGD-e *sugR*$^{F49fs}$ | This study |
| LJR248 | EGD-e *sugR*$^{D122*}$ | This study |
| LJR249 | EGD-e *sugR*$^{D122*}$ | This study |
| LJR250 | EGD-e *sugR*$^{D122*}$ | This study |
| LJR257 | EGD-e *sugR*$^{L64*}$ | This study |
| LJR258 | EGD-e *sugR*$^{F49fs}$ | This study |
| LJR259 | EGD-e *sugR*$^{F49fs}$ | This study |
| LJR260 | EGD-e *sugR*$^{F49fs}$ | This study |
| LJR261 | EGD-e ∆*fepA* | This study |
| LJR262 | EGD-e ∆*sugE1/2* | This study |
| LJR265 | EGD-e ∆*fepA* pIMK3-*fepA*; KanR | This study |

| | | |
|---|---|---|
| LJR266 | EGD-e Δ*fepA sugR*$^{D71*}$ | This study |
| LJR267 | EGD-e Δ*fepA sugR*$^{F49fs}$ | This study |
| LJR268 | EGD-e Δ*fepA sugR*$^{S44*}$ | This study |
| LJR269 | EGD-e Δ*fepA sugR*$^{A23D}$ | This study |
| LJR270 | EGD-e Δ*sugE1/2 fepR*$^{G157*}$ | This study |
| LJR271 | EGD-e Δ*sugE1/2 fepR*$^{I185fs}$ | This study |
| LJR272 | EGD-e Δ*sugE1/2 fepR*$^{D171fs}$ | This study |
| LJR273 | EGD-e Δ*sugE1/2 fepR*$^{P107L}$ | This study |
| LJR274 | EGD-e Δ*sugE1/2 fepR*$^{E89fs}$ | This study |
| LJR275 | EGD-e Δ*sugE1/2 fepR*$^{A154E}$ | This study |
| LJR276 | EGD-e Δ*sugE1/2 fepR*$^{V115fs}$ | This study |
| LJR277 | EGD-e Δ*sugE1/2 fepR*$^{V115fs}$ | This study |
| LJR280 | EGD-e Δ*fepA sugR*$^{L57*}$ | This study |
| LJR281 | EGD-e Δ*fepA sugR*$^{Y19fs}$ | This study |
| LJR282 | EGD-e Δ*fepA sugR*$^{S81*}$ | This study |
| LJR283 | EGD-e Δ*fepA sugR*$^{S81*}$ | This study |
| LJR301 | EGD-e pIMK3-*sugE1/2*; KanR | This study |
| LJR302 | EGD-e pPL3e-$P_{fepR}^{A\text{-}33G}$-*lacZ*; ErmR | This study |
| LJR303 | EGD-e pPL3e-$P_{fepR}^{G\text{-}27T}$-*lacZ*; ErmR | This study |
| LJR326 | EGD-e Δ*fepA*Δ*sugE1/2 lmo1753*$^{K19fs}$, short: Δ*fepA*Δ*sugE1/2* BAC1 | This study |
| LJR327 | EGD-e Δ*fepA*Δ*sugE1/2 lmo1753*$^{K19fs}$, short: Δ*fepA*Δ*sugE1/2* CTAB1 | This study |
| LJR328 | EGD-e Δ*fepA*Δ*sugE1/2 lmo1753*$^{K19fs}$, short: Δ*fepA*Δ*sugE1/2* CTAB2 | This study |
| LJR329 | EGD-e Δ*fepA*Δ*sugE1/2* | This study |

| | | |
|---|---|---|
| LJR330 | EGD-e $\Delta$*fepA*$\Delta$*sugE1/2 lmo1753*$^{V225fs}$ $P_{lmo1682}^{G-37A}$, short: $\Delta$*fepA*$\Delta$*sugE1/2* BAC2 | This study |
| LJR336 | EGD-e pPL3e-$P_{fepR}$-*lacZ*; ErmR | This study |

fs – frameshift

**Table S2: Primers used in this study**

| Number | Name | Sequence |
|---|---|---|
| FD1 | pWH844-*fepR* fw | AAAGGATCCAGAAAAGAAGAAATCAAACAAGCTGC |
| FD2 | pWH844-*fepR* rev | AAAGTCGACTTAATTCAAAGCTTTTAGCGTAATTCCTCTC |
| FD3 | $P_{fepR}$ fw | GACATACGAATTGATTAGCGAATTTTTAGAA |
| FD4 | $P_{fepR}$ rev | CATTCCACTCCTCTCACAAAAACTG |
| FD5 | pPL3e-$P_{fepR}$ fw | AAAGGATCCGACATACGAATTGATTAGCGAATTTTTAGA |
| FD6 | pPL3e-$P_{fepR}$ rev | AAAGTCGACTTCTTTTCTCATCATTCCACTCCTCT |
| JR247 | *sugE1/2* up fw | ACGCGTCGACGGCAGAACTAGTTAATGAGAAG |
| JR248 | *sugE1/2* up rev | TTTCAATCCGACCCCTGCCATAATCAAATAAAACC |
| JR249 | *sugE1/2* down fw | ATTATGGCAGGGGTCGGATTGAAATTAACATCTGG |
| JR250 | *sugE1/2* down rev | CGGGGTACCGGCAACTGCACCTTCTGG |
| JR262 | pIMK3-*sugE1/2* fw | CATGCCATGGGGGCTTGGTTTTATTTGATTATGGCAG |
| JR263 | pIMK3-*sugE1/2* rev | ACGCGTCGACTTAAACGCCAGATGTTAATTTCAATC |
| LMS478 | pIMK3-*fepA* fw | AAACCATGGGGGCAAAAAATATGGAAATTTTAGAAACAGATTCA |
| LMS479 | pIMK3-*fepA* rev | TTTGTCGACTTATTTAAATAAAATATGTTTTTTCTTCATATAGAATA CAATG |
| LMS484 | *fepA* up rev | TCGTTTTTATTTTGTTTCTAAAATTTCCATATTTTTTGCCATACTA |
| LMS485 | *fepA* up fw | AAAGTCGACACCAATACGTAGCAAAGATTTAGTCG |
| LMS486 | *fepA* down fw | ATTTTAGAAACAAAATAAAAACGAGACGAAGATAGATGATATC |
| LMS487 | *fepA* down rev | TTTGGTACCGCCACCTGTAAACAATAAAAGCTAAG |

**Supplementary Figures**

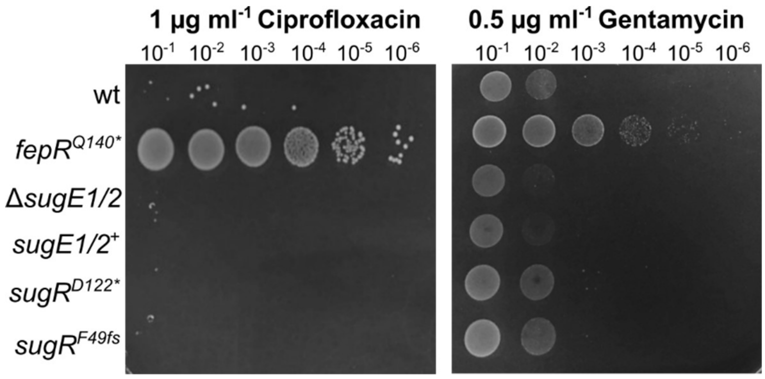


**Figure S1: Cross-resistance of *sugR* mutant strains**

Drop dilution assays of *L. monocytogenes* strains EGD-e (wt), a *sugE1/2* deletion strain (Δ*sugE1/2*), a wt strain containing the IPTG-inducible pIMK3-*sugE1/2* plasmid LJR301 (*sugE1/2*$^{+}$) and the suppressor mutants *sugR*$^{D122*}$ (LJR248), and *sugR*$^{F49fs*}$ (LJR258). The *fepR*$^{Q140*}$ suppressor mutant was used as a control. Cells were propagated on BHI plates or BHI plates containing 1 µg ml$^{-1}$ ciprofloxacin or 0.5 µg ml$^{-1}$ gentamycin. All plates were supplemented with 1 mM IPTG to induce the expression of *sugE1/2* in the *sugE1/E2*$^{+}$ strain and plates were incubated overnight at 37°C. A representative image of at least three biological replicates is shown.

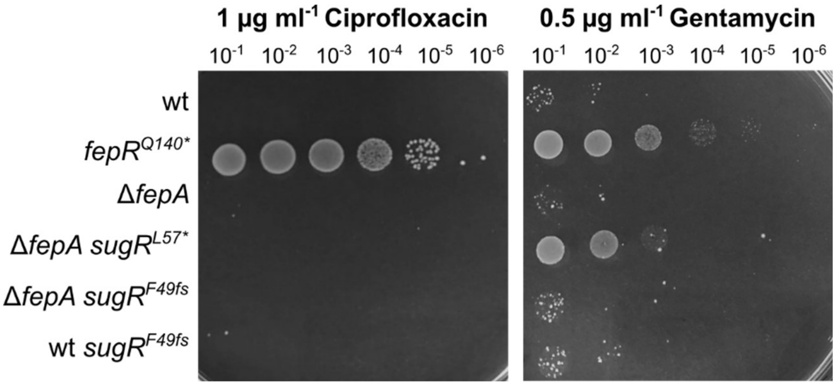


**Figure S2: Cross-resistance of Δ*fepA sugR* mutant strains**

Drop dilution assays of *L. monocytogenes* strains EGD-e (wt), the *fepA* deletion strain (Δ*fepA*) and the suppressor mutants Δ*fepA sugR*$^{L57*}$(LJR280), Δ*fepA sugR*$^{F49fs}$ (LJR267) and wt *sugR*$^{F49fs}$ (LJR258). The *fepR*$^{Q140*}$ suppressor mutant (LJR208) was used as a control. Cells were propagated on BHI plates or BHI plates containing 1 µg ml$^{-1}$ ciprofloxacin or 0.5 µg ml$^{-1}$ gentamycin and were incubated overnight at 37°C. A representative image of at least three biological replicates is shown.
